# Comprehensive structural and interactome analysis reveals novel interactions and protein binding sites in miR-675: a non-coding RNA critically involved in multiple diseases

**DOI:** 10.1101/2023.05.20.541577

**Authors:** Abhishek Dey

## Abstract

miR-675 is a microRNA expressed from exon 1 of H19 long non-coding RNA. H19 lncRNA is temporally expressed in humans and atypical expression of miR-675 has been linked with several diseases and disorders. To execute its function inside the cell, miR-675 is folded into a particular conformation which aids in its interaction with several other biological molecules. However, the exact folding dynamics of miR-675 and its complete interaction map are currently unknown. Moreover, how H19 lncRNA and miR-675 crosstalk and modulate each other’s activities is also unclear. Detailed structural analysis of miR-675 in this study determines its conformation and identifies novel protein binding sites on miR-675 which can make it an excellent therapeutic target against numerous diseases. Mapping of the interactome identified some of known and unknown interactors of miR-675 which aid in expanding our repertoire of miR-675 involved pathways in the cell. This analysis also identified some of the previously unknown and yet to be characterised proteins as probable interactors of miR-675. Structural and base pair conservation analysis between H19 lncRNA and miR-675 results in structural transformations in miR-675 thus describing the earlier unknown mechanism of interaction between these two molecules. Comprehensively, this study details the conformation of miR-675, its interacting biological partners and explains its relationship with H19 lncRNA which can be interpreted to understand the role of miR-675 in the development and progression of various diseases.

## Introduction

microRNAs (miRNAs) are small non-coding RNA which post-transcriptionally regulate gene expression by interacting with mRNA in multicellular organisms. Their interaction with mRNAs affects later stability and translation thus modulating multiple processes including proliferation, differentiation, and apoptosis etc. miR-675 is 73 nucleotides long micro-RNA which is expressed from the exon 1 of H19 long noncoding RNA (lncRNA). H19 lncRNA is temporally expressed in humans where it is expressed during the placental stage while the expression is drastically reduced after birth [1]. Interestingly, abnormal expression of H19 lncRNA has been associated with various diseases in humans including tumorigenesis, neurogenesis, angiogenesis, and fibrosis progression [1].

Being expressed from H19 lncRNA, aberrant expression of mir-675 has also been associated with various disorders in humans [2]. Enhanced expression of miR-675 is associated with indefinite proliferation and growth of hepatic cells resulting in hepatocellular carcinoma [3]. Studies have also shown that miR-675-3p (a mature form of miR-675) can promote colorectal cancer, esophageal squamous cell carcinoma and regulate skeletal muscle regeneration and differentiation [4-6]. Recently, association between abnormal expression of miR-675 and cardiovascular disease have also been identified [7]. Elevated levels of miR-675 in the patient serum suffering from atherosclerosis highlights the need to use miR-675 as a new diagnostic biomarker in early diagnosis of cardiovascular disorder [7]. Abnormal expression of miR-675 has also been linked to various neurological diseases. Downregulated expression of miR-675 has been found to be associated with the cases of treatment resistant schizophrenia (TRS) [8]. H19 lncRNA, precursor of miR-675 was also found to be highly activated in human dopaminergic neuron loss which results in Parkinson’s diseases [9]. In contrast, downregulated H19 lncRNA along with upregulated miR-129 was found to rescue PC12 cells mimicking AD model by increasing cell viability and restraining apoptosis [10].

miRNAs regulate gene expression either by repressing translation or by promoting mRNA degradation by binding to their 3’ UTR. Atypical expression of miRNAs results in an anomaly within associations between miRNA and their interactors that can result in disease progression and pathology. Multiple studies have indicated the interplay between miR-675 and multiple interactors during the advancement of various diseases. Higher plasma levels of miR-675 and its precursor H19 lncRNA in the breast cancer patients indicate an interplay between both miRNA and lncRNA [11]. Indeed, overexpression of miR-675 was able to mitigate the inhibitory effect of siRNA on H19 lncRNA in breast cancer cell line MCF-7 [11]. Interaction of mir-675-5p with serine-threonine phosphatases have been shown to activate Glycogen Synthase Kinase 3β (GSK 3β), thus enhancing the nuclear localization of β-catenin which aids in the progression of colorectal cancer [12]. Direct interaction between miR-675 and mRNA of ubiquitin 3 ligase family (c-Cbl and Cbl-b) have shown to enhance tumorigenesis and metastasis in breast cancer cells [13]. Contrary to the progression of tumours, miR-675 has also been shown to suppress tumour growth by binding to Fas-associated protein with death domain (FADD) and inhibiting its activity in cellular apoptosis, thus promoting necroptosis in human hepatocellular carcinoma cell lines [14].

Despite the advancement in identification of miR-675 interacting partners involved in the progression and metastasis of various tumours, our knowledge regarding the role of miR-675 in the advancement of other diseases is still obscure. Micro RNAs are known to fold into specific conformations which are essential for their interactions with specific proteins/interactors and hence their function. Regardless of in-depth biochemical analysis of miR-675 and its involvement in various tumours, the knowledge about its detailed folding dynamics, interacting partners and the RNA-binding protein (RBP) interacting sites on miR-675 is still missing. Understanding how the folding of miR-675 contributes its crosstalk with various proteins and RNA will be key in defining the involvement of miR-675 in the development and progression of various diseases like cancers, neurological, and cardiovascular disorders. Here we report the very first detailed conformational analysis of *in vitro* synthesised miR-675 by performing Selective 2’ Hydroxyl Acylation analysed by Primer Extension (SHAPE) assay. Multiple algorithm searches were also performed to determine the interactome network of miR-675 and to map the protein binding site on miR-675 itself. Co-fold analysis of miR-675 with its precursor H19 lncRNA was also undertaken to understand any conformational changes incurred on both interactors upon their binding that could explain the mechanistic role of miR-675 in progression of various pathogenesis or diseases. This current study identifies previously unknown interactors of miR-675. Understanding its conformation, interactome and performing co-fold analysis with H19 lncRNA further lays the foundation for future studies of miR-675 role in disease pathogenesis.

## Materials and Methods

### Selective 2’ Hydroxyl acylation analyzed by Primer Extension (SHAPE) assay

dsDNA construct of miR-675 was procured as g-blocks from Integrated DNA Technologies (IDT). Each construct is flanked by structural RNA adapters [15] and a T7 promoter region at the 5′ end. These DNA constructs were used as a template to *in vitro* transcribed RNA using T7 high yield RNA kit (New England Biolabs). The synthesized RNA was DNase treated (TURBODNase), purified using Purelink RNA mini kit (Invitrogen) and quantified with nanodrop.

Samples of 6 μg of *in vitro* transcribed RNA were denatured at 65 °C for 5 min and snap-cooled in ice. After the addition of a folding buffer (100 mM KCl, 10 mM MgCl2, 100 mM Bicine, pH 8.3), RNA was incubated at 37 °C for 10 min. The folded RNA was treated with 10 μL of 5-Nitro Isatoic Anhydride (5NIA, 25 mM final concentration). Subsequently, for negative controls (unmodified RNA) an equivalent amount of Dimethyl Sulfoxide (DMSO) was added to the folded RNA. The complete reaction mixture was further incubated for 5 min at 37 °C to allow complete modifications of the unpaired RNA nucleotides. Both the modified and unmodified RNAs were purified using the PurelinkRNA mini kit and quantified with nanodrop.

Purified RNA from above was reverse transcribed using Gene-specific reverse primer (Table S1) directed against the 3′ RNA adapter sequence and SuperScript II reverse transcriptase under error prone conditions as previously described [16]. The resultant cDNA was purified using G50 column (GE healthcare) and subjected to second strand synthesis (NEBNext Second Strand Synthesis Module). For library generation primers specific to the 5′ and 3′ RNA adapter sequence were synthesized (Table S1) and the whole cDNA was PCR amplified using the NEB Q5 HotStart polymerase (NEB). Secondary PCR was performed to introduce TrueSeq barcodes [16]. Samples were purified using the Ampure XP (Beckman Coulter) beads and the resultant libraries were quantified using Qubit dsDNA HS Assay kit (ThermoFisher). Final libraries were run on Agilent Bioanalyzer for quality check. These TrueSeq libraries were then sequenced as necessary for their desired length, primarily as paired end 2 × 151 read multiplex runs on MiSeq platform (Illumina). We used the ShapeMapper2 algorithm [17] to determine the mutation frequency in chemically modified (5NIA) and control (DMSO treated) RNA samples and to calculate chemical reactivity for each RNA nucleotide using the following equation:

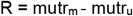

where *R* is the chemical reactivity, mutr_m_ is the mutation rate calculated for chemically modified RNA and mutr_u_ is the mutation rate calculated for untreated control RNA samples. We used this chemical reactivity to inform a minimum free energy structure using Superfold [18] and visualized the model using VARNA [19]. RNA arc and secondary structure models were generated using RNAvigate [20]. Comparative miR-675 chemical reactivity plots were generated using the plot skyline function from RNAvigate package [20]. SHAPE reactivities calculated for all the replicates and merged datasets are available in the supplementary file *miR-675_SNRNASM*.*xlsx*.

### Identification and construction of miR-675 network

To determine potential interacting partners, mapping interacting networks and to identify the protein binding sites on the miR-675, searches from three web-based algorithms: RNAinter [21], TargetScan [22] and RBPsuite [23 were executed, and each search was run on default parameters. RNAinter provides information about the interconnected partners, structure, localization, modification, experimental evidence of any queried RNA from multiple species. Scoring parameters for RNAinter were selected between 0.0 to 1.0 for RNAinter and any hits with the value less than 0.0 were excluded from further analysis.

Targetscan predicts interactors of any query miRNA by identifying the conserved sequences (8mer, 7mer and 6mer) in the 3’ UTR of potential interactors that should be complementary to the seed region of mature miRNA. Seed regions are 2-8 nucleotide long sequences in mature miRNA which are recognised by the target molecules (mRNA, miRNA, lncRNA) to form stable miRNA-target complex [24]. Predictions were segregated based on their context++ score which is a cumulative sum of multiple features like site type, supplementary pairing, local AU, minimum distance, target site abundance, seed-pairing stability etc [25].

RBPsuite is the web-based server to identify the protein binding nucleotides in any given RNA molecule. It uses two-deep learning-based algorithms to identify the RNA binding protein sites in both linear and circular RNA. Search was performed by selecting miR-675 as linear RNA/circular RNA with the general model being designated as prediction model. Protein binding site on RNA sequence is being depicted by binding score on the scale of 0-1. Higher the score, greater will be the probability of the presence of RBP site on the queried RNA sequence. RNA motifs less than 0.5 binding scores were excluded. If the server identifies a verified motif with binding score more than 0.5, then those are highlighted in red, and the motif logo is provided in the output.

### RNA co-fold analysis

Co-fold analysis between miR-675 and H19 lncRNA exons was performed using RNAcofold algorithm from vienna RNA websuite [26, 27]. RNAcofold calculates hybridization energy and base pairing probabilities between two interacting RNA molecules. RNAcofold links the two sequences together and the junction is treated as an exterior loop while calculating the hybridization energies. In addition to the minimum free energy structure, the algorithm also provides the fraction of suboptimal structures and the equilibrium concentrations of the duplexes formed. Base pairing conservation was computed to identify the changes in the base pairing pattern within two interacting RNA molecules.

### Statistical analysis

Most of the statistical analyses were performed using the R software package unless otherwise stated. Pearson correlation test was used to evaluate correlation between chemically probed replicates (*n* = 2) across all test samples. Validity of the folding conformation of two replicates was accessed by the scorer algorithm embedded in RNAstructure package [28]. Scorer determines the Positive predictive value (PPV) and sensitivity (Sens) between two RNA conformations. Positive predictive value (PPV) is the fraction/percentage of the base pairs in the predicted minimum free energy (MFE) structure where the known structure is determined by comparative sequence analysis. Sensitivity (Sens) is the maximum expected accuracy of RNA structure determined by fraction/percentage of known base pairs correctly predicted in minimum free energy (MFE) structure where the known base pairs are determined by comparative sequence analysis.

## Results

### Secondary structure of miR-675 represents canonical stem loop helical conformation

To experimentally analyse the folding conformation of nascent miR-675, mutational profiling was performed after treating synthetic miR-675 with 5NIA. Being a SHAPE reagent, 5NIA is known to modify the backbone of RNA molecules by forming a chemical adduct with 2’ hydroxyl group of the unpaired RNA nucleotides [29]. The modifications can be identified by mutational profiling followed by massively parallel sequencing. SHAPE analysis of the two independent replicates (n=2) of *in vitro* transcribed miR-675 resulted in chemical reactivity profile for both replicates with most of the nucleotides having low chemical modification except for nucleotides G12, U13, A18, A34, C35, U36, U37, G38, G39, U40 and A49 (Fig. 1A, 1B). This suggest that these nucleotides are highly flexible and are not involved in base-pairing interactions. Comparative analysis showed that both replicates have identical chemical reactivity profiles (Pearson Correlation coefficient, R= 0.98) (Fig. 1C, 1D) while Scorer analysis [30] on the models obtained from replicates showed that both RNA folds into an identical conformation with the positive predictive value (PPV) and sensitivity (Sens) of 100% (Fig. 1E). Once it was confirmed that both replicates’ folds into an identical conformation, average nucleotide chemical profiling was generated (by merging the fastq files for both replicates and re-running the shapemapper algorithm on the merged fastq files) and the final model was created for miR-675 (Fig. 1H).

**Figure 1:**
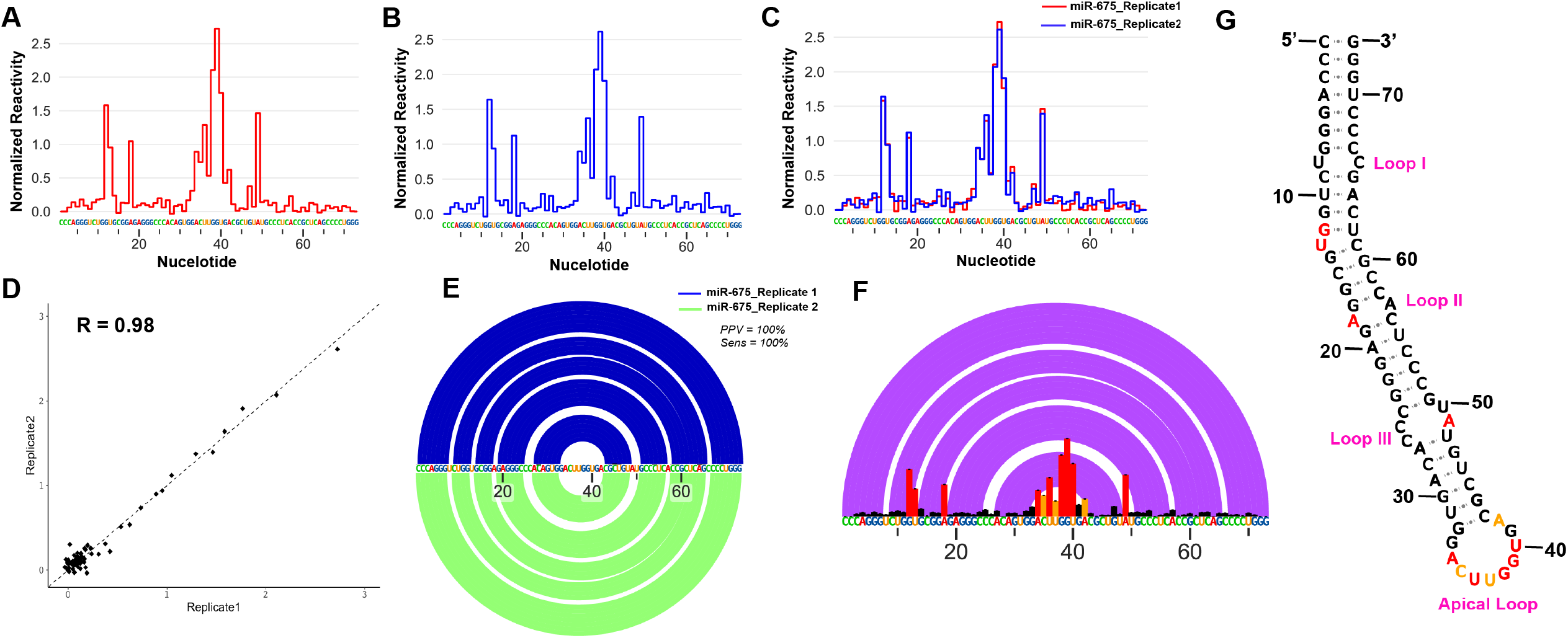
*In vitro* architecture and statistical analysis of miR-675 **A**. Normalized 5-NIA reactivity profile for *in vitro* synthesized miR-675 replicate 1 **B**. Normalized 5-NIA reactivity profile for *in vitro* synthesized miR-675 replicate 2. Some of the nucleotides were found to have high preference for 5-NIA as compared to others in both the replicates. **C**. Overlapped normalized 5-NIA reactivity profile of *in vitro* synthesized miR-675 replicates. miR-675-Replicate 1 is represented in red color while miR-675-Replicate 2 is represented as blue color. The reactivity profile was found to be highly similar for both the replicates. **D**. Scatter plot with Pearson Correlation coefficient between the normalized reactivity profiles of miR-675 replicates. R=0.98 illustrates identical and reproducible reactivity profiles between both the replicates of miR-675. **E**. Overlapped RNA arcplot (depicting secondary structure) with scorer results for miR-675, Replicate 1 and Replicate 2. Both miR-675 replicates fold into identical conformation with PPV=100% and SENS=100%. **F**. Normalized 5-NIA reactivity profile generated after averaging the reactivity profiles of miR-675 replicate 1 and replicate 2. Also shown is the overlapped arc profile representing the secondary structure of miR-675. **G**. Secondary structure model of miR-675 was obtained after feeding superfold with obtained SHAPE constraints. Black color represents 5-NIA reactivity <0.4, orange represents 5-NIA reactivity between 0.4 and 0.8 while Red represents 5-NIA reactivity >0.8 for nucleotides.

Based on the SHAPE constraints and minimum free energy, miR-675 folds into a stem loop helical conformation which includes multiple internal bulges or loops and culminates in an apical loop. The basal stem of the miR-675 helix was stabilised by 7 base pairs (nts 1-7 and nts 67-70) which includes 6 GC base pairs (Fig. 1H, 1G). There are three internal loops (Loop 1, Loop 2 and Loop 3) and a single bulge (Bulge I). Loop I and Loop II are formed by 2 unpaired or flexible nucleotides (U8 and C67-Loop 1) and (A18 and A57-Loop 2) (Fig. 1G) while Loop III was the largest internal loop which was formed by 4 unpaired nucleotides (C25, C26, A49, U50) in the stem loop helix structure of miR-675. A single nucleotide bulge (U13) was also present which was sandwiched between internal loop 1 and loop 2. Interestingly, only nucleotides U13 (bulge I), A18 (loop II) and A49 (loop III) were found to be accessible to SHAPE reagent, 5NIA, thus suggesting that the remaining inaccessible nucleotides present in loops were either arranged alternatively or involved in intermolecular interaction thus resulting in minor conformations which are energetically least favourable and hence remain undetected in current study. The apical end of the miR-675 ends with a formation of 10 nucleotide apical loop starting from G33 to A42 (5’-GACUUGGUGA-3’) (Fig. 1G). Closer examination of the loop nucleotides suggests that most of them were highly reactive to 5NIA, thus suggesting that they were not involved in any base-pair interactions and were highly flexible. A hexanucleotide base pairs forms a bridge between internal loop 3 and terminal apical loop (Fig. 1G). Overall, the chemical probing strategy of miR-675 reveals a tight canonical stem-loop helical conformations with most nucleotides involved in base-pairing interactions thus being inaccessible to 5NIA.

### miR-675 interacts with array of biological molecules

microRNAs are known to regulate gene expression by interacting with various biomolecules inside the cells. Any abnormalities in this interaction will result in altered gene expression and disease conditions. Multiple studies have identified various interactors of miR-675, however the overall interaction network of miR-675 is still obscure. To identify the molecular partners of miR-675 and to map its interactome multiple searches with three different algorithm (RNAinter, RBPsuite and TargetScan) were performed. Search with RNAinter resulted in approximately 300 interactors with varying scores with miR-675. However, interactors with the binding score between 1.0-0.0 were taken forward for further analysis. This segregation resulted in the identification of 168 interactors (Fig. 2A) which mainly belong to three categories viz. Transcription factor (86.8%), Histone modifications (9.5%) and Proteins (4.7%) (Fig. 2B). Although the total number of histone modifications identified as possible interactors of miR-675 is very less than the transcription factor, the top 5 miR-675 interactors identified from RNAinter belong to the group of former with the binding score ranging between 0.77 to 0.72 (Fig. 2A, Table S2). Nevertheless, results from RNAinter suggest that miR-675 is involved in regulating gene expression by interacting with epigenetic regulators and various transcription factors.

**Figure 2:**
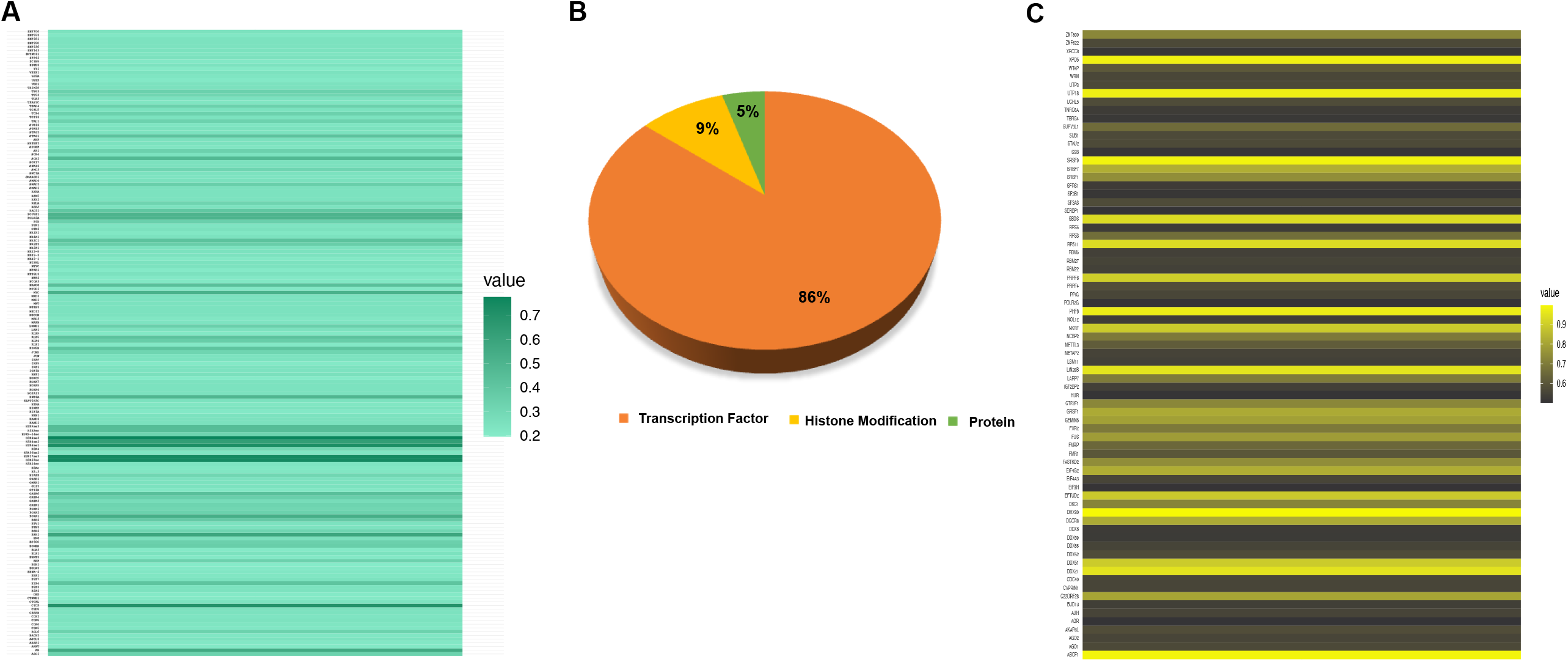
miR-675 interacting partners identified from RNAinter and RBPsuite. **A**. Heatmap of miR-675 interacting partners identified from RNAinter. The interaction score ranges from 0.8 to 0.2, with 0.8 means strong interactions and 0.2 means weak interaction with miR-675. **B**. Pie chart depicting the percentage of miR-675 interactors identified through RNAinter. ∼86% of miR-675 interacting partners belong to the group of transcription factors. Interestingly, 9% of histone modification sites were also identified as possible interaction sites for miR-675 and the remaining 5% belong to the group of other proteins which can also interact with miR-675. **C**. Heatmap of RNA binding proteins from RBPsuite bound to miR-675. The binding scores for these proteins range from ≥ 0.9 to ≤0.6, with ≥ 0.9 means strong binding and ≤0.6 means weak binding.

Within a cellular system, RNA Binding Proteins (RBPs) are involved in multiple biological processes including gene expression and regulation of cellular pathways. Determining the RBPs binding site on RNA is essential to gain mechanistic understanding on above processes which are regulated through microRNA or non-coding RNA. RBPsuite was used to determine the miR-675 binding proteins and its motif/sequences involved in RNA-protein interactions. The results were segregated based on the binding scores (Fig. 2C) and only proteins having binding score more than 0.5 are reported here (Table S3). Proteins with verified motif for miR-675 are also reported with sequence logo (Fig. 3). The obtained sequence was further mapped on the stem-helix loop model of miR-675 (Fig. 3). Interestingly, most of the motifs were found to be assembled from nucleotides which either constitute bulge, internal loop, or the apical loop of miR-675. FUS protein motif was found to be located on the bulge I of miR-675 (Fig. 3A, 3G). Surprisingly, for proteins FXR2, Lin28B and SRSF1, RBPsuite identified a common motif spanning from G16-G23 for FXR2 and Lin28B (Fig. 3) and A18-C26 for SRSF1 (Fig. 3). Nevertheless, this region also contains nucleotide A18 which is the part of loop II in the stem loop helix model of miR-675. Motif for SRSF9 was detected at the downstream end of SRSF1 motif spanning from A27-A35, which also includes a part of the apical loop (Fig. 3). Apical loop of miR-675 was also found to harbour the binding site for Human Antigen R (HuR) protein (Fig. 3F, 3G).

**Figure 3:**
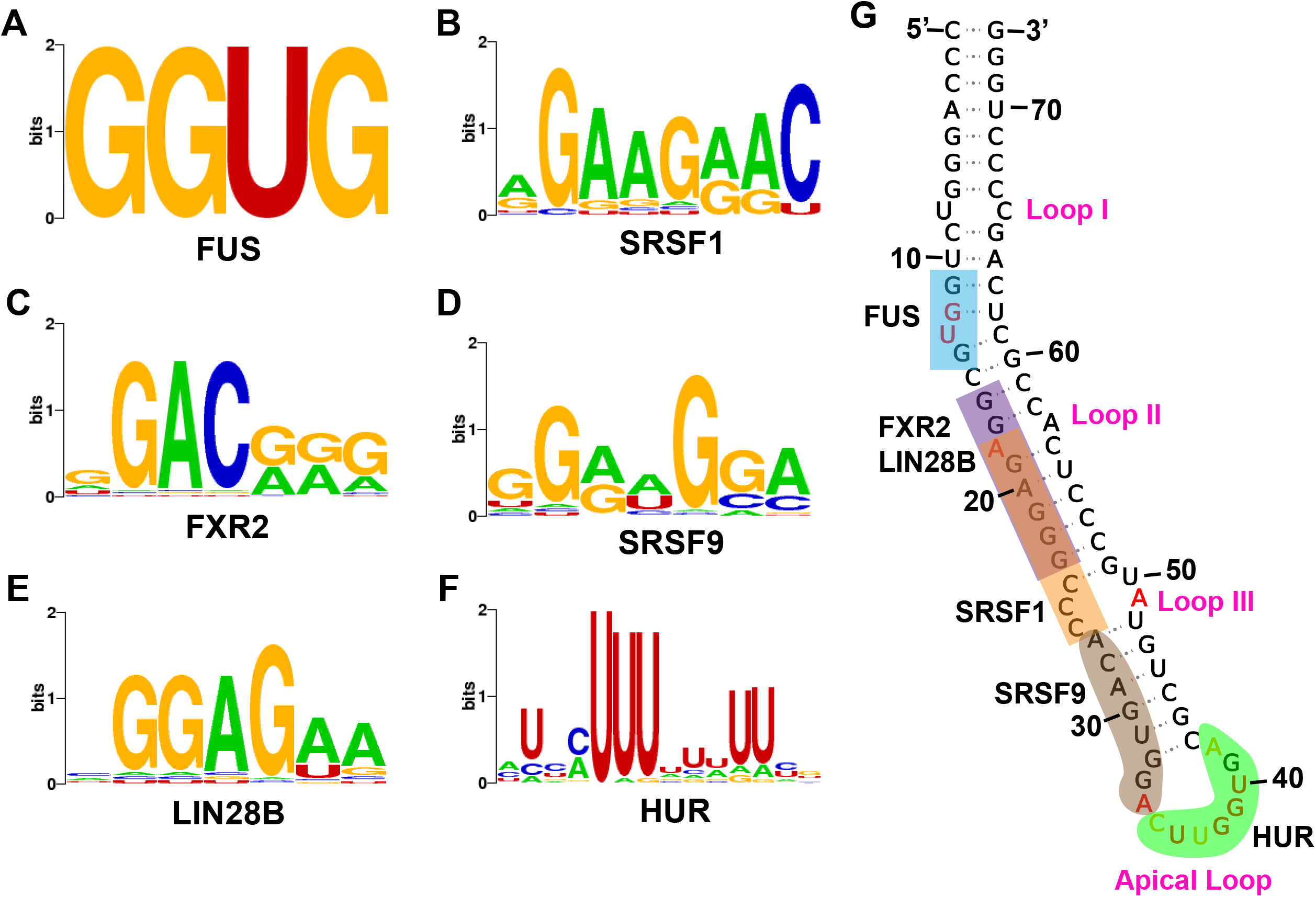
miR-675 protein binding sites of different RNA binding proteins identified through RBPsuite. miR-675 binding sequences identified for proteins **A**. Fused in Sarcoma (FUS) **B**. Serine/arginine-rich splicing factor 1 (SRSF1) **C**. FMR1 autosomal Homolog 2 (FXR2) **D**. Serine/arginine-rich splicing factor 9 (SRSF9) **E**. Lin-28 homolog B (Lin28B) **F**. Human Antigen R (HuR) **G**. Protein binding sites in miR-675. Most of the protein binding sites in miR-675 include the nucleotides from the loop region thus making miR-675 an excellent target of therapeutic importance.

To further identify additional targets of miR-675, targetSCAN analysis was performed. Total 2599 hits were identified by targetSCAN as potential interactors of miR-675. Since microRNAs execute gene regulation by binding to the 3’ untranslated regions (3’ UTRs) of mRNA, hits from targetSCAN were further sequestered based on the number of binding sites in 3’UTR of each identified interactor (Fig. 4A). Most of the interactors identified have more than one 8 nucleotides seed binding site for miR-675 (Fig. 4A). Furthermore, some hits also displayed 7 nucleotide and 6 nucleotide seed binding sites for miR-675 (Fig. 4A). In addition to the seed binding site for each microRNA, targetSCAN also relies other parameters to partition the predictions or hits. Cumulatively, these features give rise to total context++ score and cumulative weighted context++ score. Score ≤0.4 indicates better binding interactions between miRNA and its target molecule. Analysis of miR-675 cumulative weighted context++ score shows that most of the predicted interactors have scored less than 0.4 thus indicating higher confidence of interactions between the two molecules (Fig. 4B). Since targetSCAN resulted in around 2599 probable interactors for miR-675, these interactors were further categorised into different families/groups of protein (Fig. 4C). Protein families which have ≤4 members were annotated as others (30%) (Fig. 4C). The largest member of protein families which probably act as an interactor to miR-675 was found to be various receptors (∼8%), Zinc finger proteins (ZNF) (∼5%), transferase and kinase (∼3%) (Fig. 4C). ZNF proteins are implicated in numerous cellular processes like transcriptional regulation, cell migration and DNA repair pathways [31] while receptors, kinases and transferases are majorly involved in cell signalling and cellular transduction [32, 33].

**Figure 4:**
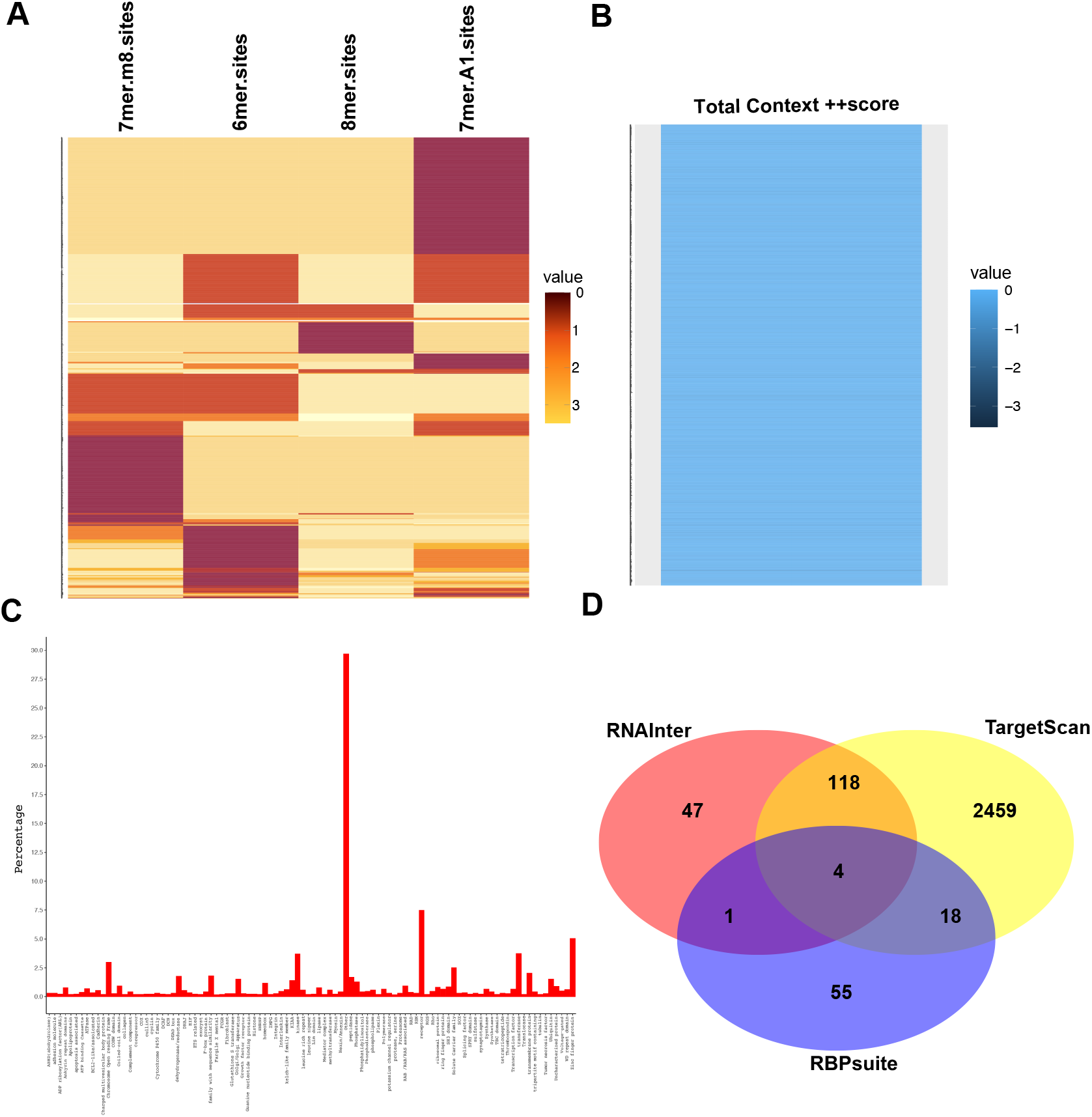
miR-675 interacting partners identified from targetSCAN. targetSCAN identifies the partners based on the miRNA target binding sites in mRNA. **A**. Heatmap of targetSCAN identified interactors with miR-675 based on the number of miRNA binding sites present in the 3’ UTR of target mRNA. **B**. Heatmap of miR-675 identified interactors from targetSCAN based on their total context++ score of binding. **C**. Bar Graph depicting the fractions of each protein family identified by targetSCAN. As stated in the main text protein families ≤4 members were annotated as others (30%). This is followed by proteins belonging to the families of receptors, kinases, and transferases. **D**. Venn diagram highlighting the number of common and unique interactors identified from RNAinter, RBPSuite and targetSCAN. As stated earlier 118 indicators were common to RNAInter and targetSCAN, 18 interactors were common to RBPsuite and targetSCAN and only 1 common interactor was identified between RBPsuite and RNAinter. About 4 interactors were commonly identified from RNAinter, RBPsuite and targetSCAN.

Comparative analysis between the interactors identified from all the three searches (RNAinter, RBPsuite and targetSCAN) resulted in the identification of common interactors (Fig. 4D). Four interactors (Zinc Finger protein (ZNF), Argonaute (AGO) protein, Eukaryotic initiation factors (EIFs) and DNA directed RNA polymerase (ploR)) were commonly identified by all the three, while 118 indicators were common to RNAInter and targetSCAN, 18 interactors were common to RBPsuite and targetSCAN and only 1 common interactor was identified by RBPsuite and RNAinter (Fig. 4D). Overall, the search resulted in an array of miR-675 interactors to which it can either bind directly or to the 3’UTR of their mRNA and can modulate gene expression and cellular processes.

### H19 long noncoding RNA (lncRNA) interacts with miR-675 by modifying its conformation

Although miRNAs regulate various cellular activities by interacting with a wide array of biological molecules, their activities are themselves controlled by the sponging effect of lncRNA, where lncRNAs can interact directly with miRNA and sequester them from their respective pathways. It has been known for a while that H19 lncRNA can interact with miR-675 [11], however how H19 lncRNA and miR-675 crosstalk is currently unknown. To understand the nature of interactions between miR-675 and H19 lncRNA, RNAcofold followed by comparative base-pair conservation analysis between individual RNA and co-folded hybrid molecules was performed. Surprisingly, it was found that all the H19 exons interact with miR-675 and these interactions rearrange the intramolecular base-pair interactions within both H19 lncRNA and miR-675 itself (Fig. 5). Interactions between H19-exon1 and miR-675 resulted in two identical heterodimeric conformations with different binding energies (Table S4). Closer examination of both models suggests that the 5’ end of miR-675 interacts with the 3’ end of H19-exon1 with global rearrangements in nucleotides interactions and opening of miR-675 stem loop helical conformation (Fig. 5A). Similarly, interactions between H19-exon3 and miR-675 also resulted in two identical heterodimeric conformations with global rearrangements in nucleotide pairing (Fig. 5C). This rearrangement also resulted in the opening of miR-675 conformation (Fig. 5C) and binding of 5’ end of miR-675 to 3’ end of H19-exon3 lncRNA.

**Figure 5:**
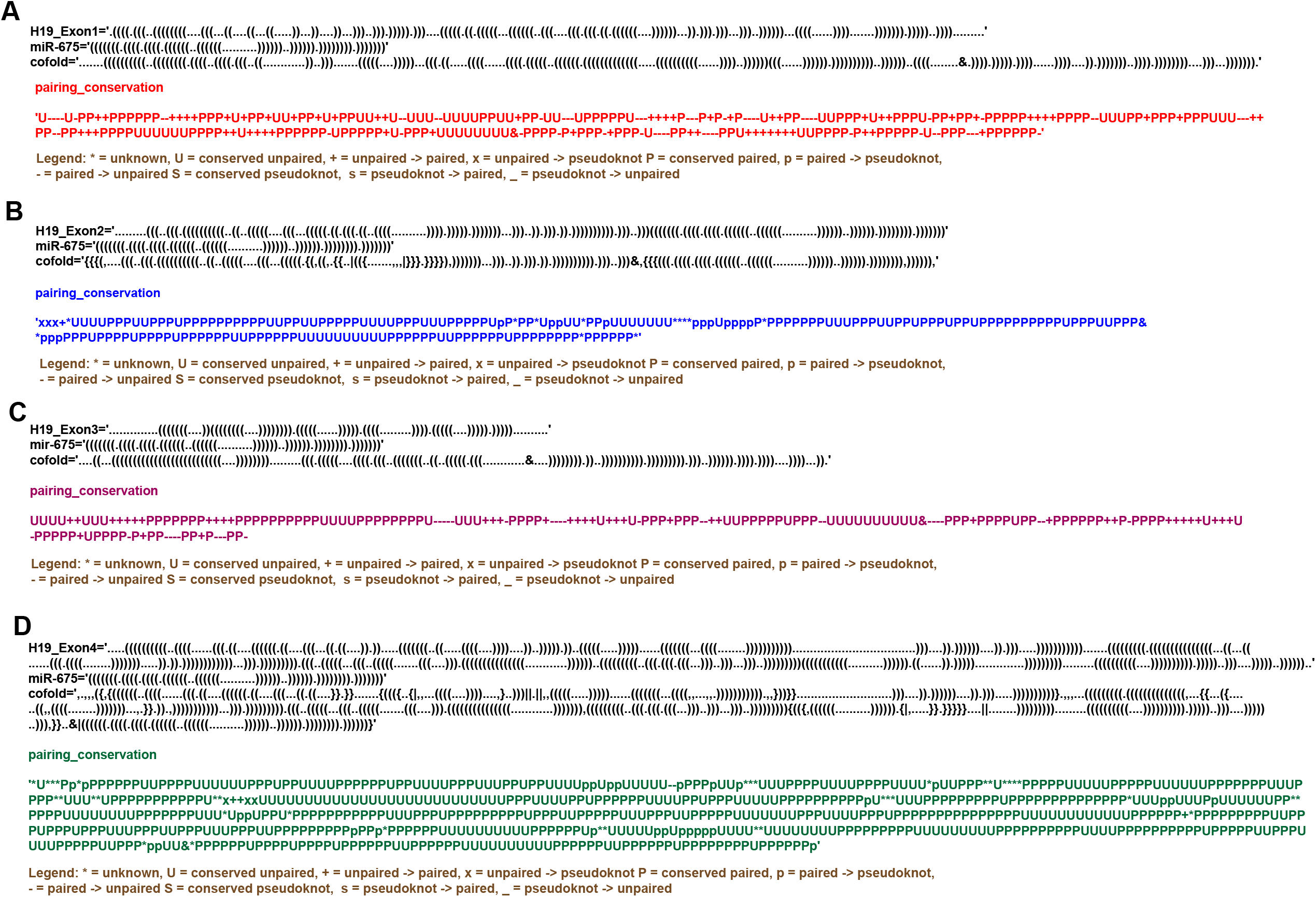
RNA cofold and base pair conservation analysis between miR-675 and exons **A**. Exon 1 (Model 1) **B**. Exon 2 (Model 2) **C**. Exon 3 (Model 1) **D**. Exon 4 (Model 2) of precursor H19 lncRNA. Upper panel in each figure represents the folding dynamics of individual H19-Exons, miR-675 and their co-folding pattern analysed by RNAcofold [26, 27] in dot-bracket notation. “&” in co-fold pattern of hybrid structure is the separator between two RNA molecules. Middle panel represents the H19-Exons and miR-675 base pairs which either got modified or remained unchanged during co-folding of both the RNA molecules. Lower panel explains the meaning of each symbol/notations used to determine the base-pair conservation during the co-folding process of H19-exons and miR-675. miR-675 had shown to prefer Exon 1 and Exon 3 of H19 lncRNA resulting in opening of its canonical stem-loop helical structure while its interaction with Exon 2 and Exon 4 results more in localized rearrangements in base-pairing between nucleotides.

However, interactions between H19-exon2/exon4 with miR-675 resulted in two different heterodimeric conformations with different binding energies (Table S4). One model consists of unperturbed folding dynamics of both H19-exon2/exon4 lncRNA and miR-675 thus suggesting that miR-675 doesn’t interact with the exon2 and exon4 region of H19 lncRNA. Interestingly, the second model however resulted in some localised re-arrangements in the base pairing patterns of both H19 lncRNA and miR-675 (Fig. 5B, 5D). This demonstrates occurrence of minute and subtle base pair rearrangements upon the interaction between exon2/exon4 of H19 lncRNA and miR-675. This interaction however does not result in the destabilisation of miR-675 helical conformation (Fig. 5B, 5D).

## Discussion

microRNAs (miRNAs) are known to regulate gene expression and other cellular pathways by influencing translation and stability of mRNAs. To execute these miRNAs had to adopt a stable conformation which aids in its interaction with other biomolecules inside the cell. Disruption in microRNA expression/activity results in the development of anomalies like tumours and neurological disorders. Determining the structural complexity and interacting partners of any miRNAs will be pivotal in understanding the pathways they are involved in and will further help in determining the progression of these disorders. The current study reports the very first secondary structure model of miR-675 and its unknown interactors. With RNAcofold and base pair conservation analysis the mechanism of interaction between H19 lncRNA and miR-675 was also elucidated for the very first time.

The current miR-675 model folds into a canonical stem loop helical conformation consisting of 3 internal loops and a bulge which are sandwiched between a basal stem and a terminal apical loop (Fig. 1H). The basal stem is highly stabilised due to base pairing interactions between 6 GC nucleotides. Most of the miR-675 was chemically unmodified thus suggesting highly stable conformation (Fig. 1H). However, nucleotides comprising loops (especially apical loop nucleotides) and bulges were found to be highly reactive to SHAPE reagent thus illustrating that they are flexible and single stranded in the final model (Fig. 1H). Within the cells these loops/bulges act as the protein binding sites and thus form the basis of RNA-protein interactions. It is possible that nucleotides involved in the formation of internal and apical loops of miR-675 act as the site for RNA-protein interactions. Indeed, analysis from RBPsuite identified proteins whose binding sites involved nucleotides of loop II, loop III, and apical loop of miR-675. More in-cell RNA SHAPE analysis will be required to accurately pinpoint the different protein binding sites on miR-675. Repeated attempts to understand the folding dynamics of the mature form of miR-675 (miR-675-3p and miR-675-5p) resulted in failure. It seems that the mature forms of miR-675 adopt more single stranded nature within cells and they execute their regulatory effect by identifying their complementary sequences in the 3’UTR of mRNA.

To accurately map miR-675 interactome and identify its binding partners, three web-based tools were searched simultaneously. Search with RNAinter resulted in the identification of various transcription factors and histone modification sites as the possible interactors of miR-675. Histone modifications are the basis of epigenetic regulation and interactions between these regions and miR-675 suggest mutual epigenetic role between the two. Furthermore, these interactions suggest that miR-675 is involved in regulating certain pathways that are implicated in the onset and development of various disorders by controlling the extent of histone methylation/acetylation. Indeed, it was found that during hepatocarcinogenesis, miR-675 epigenetically regulates histone modification sites like H3K9me3, H3k27me3 resulting in disease development and progression [34]. Additionally, RNAinter search also identified multiple transcription factors (∼87%) as possible interactors for miR-675. CTCF and ESR1 were one of the transcription factors which showed stronger interaction with miR-675 (0.69 and 0.58 respectively, Table S2). CTCF is a CCCTC-binding factor which regulates the expression of H19 lncRNA and miR-675 in particular [35]. It has been known that CTCF/h19/miR-675/Histone deacetylase (HDAC) pathway results in the differentiation of bone marrow mesenchymal stem cells into adipocytes [36]. However, the exact mechanistic role of CTCF/h19/miR-675/Histone deacetylase axis in the hepatic cancer is still not clear and will further require more in-depth biochemical analysis to understand the same. Estrogen Receptor 1 (ESR1) is a key ligand activated transcription factor for normal growth, metabolism, and sexual development. Role of ESR1 has also been implicated in the various disorders like paediatric asthma [37], breast cancer [38], and Alzheimer’s disease [39]. Elevated levels of miR-675 was found in the children’s suffering from asthma [37] however, the exact correlation between miR-675 and ESR1 in the progression of breast cancer and Alzheimer’s disease is still not clear though ESR1 is known to interact with various other miRNA during these diseases [38, 39]. It would be interesting to further investigate the role of miR-675-ESR1 axis in the onset and development of breast cancer and Alzheimer’s disease.

miRNAs generally interact with the 3’UTR of their targeted mRNA and confer gene regulation accordingly. However, there are instances where miRNAs also bind to specific RNA binding proteins (RBP) [40, 41]. These interactions are not only useful in the miRNA biogenesis but also have been implicated in the development of several diseases especially tumours, melanomas, and neurological disorders [41]. Search with RBPsuite not only resulted in identification of additional protein interactors but also determined the protein binding footprints on miR-675. Amongst all, binding sites for FUS, SRSF1, SRSF9, FXR2, LIN28B and HUR proteins were clearly identified on miR-675. Fused in Sarcoma (FUS) protein, an RNA binding protein is known to aid in gene silencing by simultaneously interacting with both miRNA and mRNA [42] and impairment in FUS protein has also been linked with several neurodegenerative diseases like amyotrophic lateral sclerosis (ALS) [42]. Lin-28 homolog B (Lin28B) is an evolutionary conserved RNA binding protein which can bind to precursor miRNAs and can interfere with their maturation process [43]. Overexpression of Lin28b is known to be linked with the development of ovarian cancer [43] and cellular proliferation in acute myeloid leukaemia [44]. Likewise, HuR is an RNA binding protein belonging to the embryonic lethal and altered vision (ELAV) family of proteins [45] and is essential for the stability and translation of mRNAs [40]. In this study, it was observed that HuR may bind to the apical loop of miR-675 (Fig. 3G). Though it primarily binds to mRNA, interactions between HuR and other miRNA have also been linked to various disorders. For instance, interaction between HuR and let-7a miRNA represses the expression of c-myc, thus advancing tumorigenesis in cervical cancer cell lines [46]. In another case inhibition of miR-16 and miR-331 by HuR resulted in the promotion and development of colorectal and prostate cancer [47, 48]. It is possible that the binding of HuR to miR-675 might have some kind of regulatory impact (positive or negative) on either of the interactors thus resulting in disease progression. As stated above these proteins interact with the nucleotides which are present in the loops and bulges of miR-675 (Fig. 3G). These single-stranded regions within miRNAs have therapeutic importance and serve as an excellent site for designing small molecule inhibitors and Antisense Oligos (ASOs). Small molecules tend to bind to these small pockets or grooves of miRNAs and can impact their biological activity while interaction between ASOs and miRNA can destabilise miRNA leading to its degradation inside the cell. Targeting miR-675 with either small molecule inhibitor or ASOs could be a way to develop novel therapeutic intervention against certain diseases like cancers and neurodegenerative disorders. Similarly, the protein binding site of miR-675 can be targeted with small molecules or ASOs to develop novel therapeutic platforms to combat above diseases.

Cellular interactions and cell signalling pathways play an indispensable role in tumour progression and metastasis. Cancerous cells interact with each other thus promoting cellular proliferation and their survival rate. Identification of cellular receptors by targetSCAN as potential interactors of miR-675 suggest that miR-675 can develop and aid in the progression of tumours and malignancies by regulating the expression of several receptors on the cell surface. Elevated levels of miR-675 have been found to upregulate the expression of tyrosine kinase receptors thus resulting in contentious division of breast cancer cell lines [13]. Other prominent interactors identified for miR-675 from targetSCAN searches were ZNF protein. ZNF can act as transcriptional regulators and regulating their expression can give rise to various tumours and malignancies. Many microRNAs are known to regulate ZNF protein levels which in turn control disease progression. Reduced level of miR-491-5p results in increased level of ZNF-703 which results in Breast Cancer Progression [49]. Surprisingly, search with targetSCAN also resulted in the identification of 23 uncharacterized proteins as potential interactors of miR-675 (Fig. 4C). Further biochemical and biophysical studies will be required to characterize these proteins, investigate the extent of their interactions with miR-675, and to determine their biological consequences.

Deeper analysis on the results obtained from the above searches (RNAInter, RBPsuite, targetSCAN) resulted in the identification of 4 common miR-675 interacting partners: ZNF, Argonaute (AGO), Eukaryotic Initiation Factors (EIF) and DNA directed RNA polymerase (polR). Identification of AGO was not surprising as it is an essential protein involved in RNA induced silencing complex (RISC) mediating gene regulation through RNA interference [50]. Though, exact mechanistic role of miR-675 and EIF is unknown, it has been found that miR-141 induced during viral infection targets EIF4E mRNA resulting in shut down of host translation machinery and promoting viral proliferation [51]. Currently, it is unknown how miR-675 expression and activity is regulated during viral infection like Hepatitis C virus (HCV), Japanese encephalitis virus (JEV), severe acute respiratory syndrome-Coronavirus 2 (SARS-CoV 2) or other viral infections. However, more studies are required to determine the unique effect of miR-675 during these viral infections which will aid in proliferation of virus and its pathogenesis, thus making miR-675 a novel anti-viral therapeutic target against above diseases.

Though being expressed from the exon 1 of H19 lncRNA itself, both miR-675 and H19 lncRNA interact to modulate each other activities thus exerting feedback regulatory mechanisms. As stated above, interactions between miR-675 and its parent molecule H19 lncRNA has been implicated in many disorders however the nature and extent of these interactions are still unknown. Here in conjunction with RNAcofold and base-pair conservation analysis, we have shown for the first time the detailed co-interaction pattern between individual exons of H19 lncRNA and miR-675. Interaction between miR-675 and exon1/exon3 of H19 lncRNA resulted in the opening of miR-675 stem helical structure as the 5’ end of latter interacts with the 3’ end of former suggesting of global rearrangements in miR-675 (Fig. 5A, 5C). Interactions between miR-675 and exon 2/exon4 however resulted in very localized subtle nucleotide rearrangements in miR-675 (Fig 5B, 5D). This suggests that miR-675 preferably binds to exon 1 and exon 3 of H19 lncRNA. Exact consequences of these interactions are yet to be determined; however, one hypothesis is that binding of miRNA to lncRNA results in the degradation of lncRNA by mimicking the targets of miRNA [52]. Alternatively, binding of H19 lncRNA and opening of miR-675 helical conformation also suggests that H19 lncRNA is sequestering away miR-675 from its intended target and thus modulating the gene expression. It will be interesting to see whether interactions between miR-675 and H19 lncRNA upregulate or downregulate the activity of either of them or both. More structural, biochemical, and cell - based reporter assays will be required to ascertain the biological consequences of this interaction and how they shape up disease progression.

Comprehensively, the work described here provides the first detailed analysis of miR-675 stem-loop helical conformation. Based on the secondary structure model of miR-675 and interaction analysis, this study allows identification of previously unknown protein binding sites which can be further harnessed to develop small molecule inhibitors or ASOs against miR-675. Interactome analysis also determines some of the very first unknown and uncharacterised interactors of miR-675. Detailed analysis between H19 lncRNA and miR-675 illustrates the preference of miR-675 for exon 1 and exon 3 of H19 lncRNA. Opening of miR-675 is indicative of the sponging effect of H19 lncRNA over miR-675. This contributes to the necessary groundwork for future mutational and genetic analyses and will further aid in detailed understanding of the mechanism of miR-675 biogenesis, and its function in various disorders and pathogenic stages.

## Supporting information

miR-75_SNRNASM

## Funding

This work is supported by the Ramalingaswami Re-entry fellowship awarded to Dr. Abhishek Dey by the Department of Biotechnology-India.

### This article bears the NIPER-R communication number

NIPER-R/Communication/430

## Data Availability

The author confirms that the data supporting the findings of this study are available within the article and its supplementary materials.

## Supplementary Figure

**Figure S1:**

RNA cofold and base pair conservation analysis between miR-675 and exons **A**. Exon 1 (Model 2) **B**. Exon 2 (Model 1) **C**. Exon 3 (Model 2) **D**. Exon 4 (Model 1) of precursor H19 lncRNA. Upper panel in each figure represents the folding dynamics of individual H19-Exons, miR-675 and their co-folding pattern of hybrid structure analysed by RNAcofold [26, 27] in dot-bracket notation. “&” in co-fold pattern of hybrid structure is the separator between two RNA molecules. Middle panel represents the H19-Exons and miR-675 base pairs which either got modified or remained unchanged during co-folding of both the RNA molecules. Lower panel explains the meaning of each symbol/notations used to determine the base-pair conservation during the co-folding process of H19-exons and miR-675.

## Supplementary Table

**Table S1:**

Primers Used for the SuperScript II Error Prone Reverse Transcriptase PCR and Library Generation.

**Table S2:**

List of miR-675 interactors identified from RNAinter.

**Table S3:**

List of RNA binding proteins binding to miR-675 identified from RBPsuite.

**Table S4:**

Free binding energies of H19 lncRNA and miR-675 heterodimers identified from RNAcofold.

