## Supplemental Figure 1 for "Comprehensive structural and interactome analysis reveals novel interactions and protein binding sites in miR-675: a non-coding RNA critically involved in multiple diseases"

**A**

H19_Exon1='.((((.(((..((((((((....(((...((....((...((.....))...))....))...)))..))).))))).)))....(((((.((.(((((...((((((..(((....(((.(((.((.((((((....))))))...)).))).)))...)))..))))))...((((......))))........))))))).)))))..))))........'
miR-675='(((((((.((((.((((.((((((..((((((..........))))))..)))))).)))))))).)))))))'
cofold='......,((((((((((..((((((((.((((..((((.{((..((...,,...,...))..}))........(((((....)))))...{((.((.....((((.......((((.(((((..((((((.(((((((((({(({,...((((((((((......))))..))))))(((......)))))).))))))))))..))))))..((((.........&.)))).))))).))))......))))....)).)))))))..)))).))))))))).....)))...))))))),'

pairing_conservation

'U----U*PP++PPPPPP--++++PPP+U+PP+UU+PP+UxPPUU++U--**U--*UUUPPUUxPP-UU---UPPPPPU---++++P---p+P-+P----U++PP----UUPPP+U++PPPU-PP+PP+-PPPPP++++PpPPp*UUUPP+PPP+PPPUUU---++PP--PP+++PPPPUUUUUUPPPP++U++++PPPPPP-UPPPPP+U-PPP+UUUUUUUU&-PPPP-P+PPP-+PPP-U----PP++----PPU+++++++UUPPPP-P++PPPPP-U--PPP---+PPPPPP*'

Legend: * = unknown, U = conserved unpaired, + = unpaired -> paired, x = unpaired -> pseudoknot P = conserved paired, p = paired -> pseudoknot,
- = paired -> unpaired S = conserved pseudoknot, s = pseudoknot -> paired, _ = pseudoknot -> unpaired

**B**

H19_Exon2='.........(((..(((.((((((((((..((..(((((....(((...(((((.((.(((.((..((((..........)))).))))).)))))))...)))..)).))).)).)))))))))).))).))(((((((.((((.((((.((((((..((((((..........))))))..)))))).)))))))).)))))))'
miR-675='(((((((.((((.((((.((((((..((((((..........))))))..)))))).)))))))).)))))))'
cofold='.........(((..(((.((((((((((..((..(((((....(((...(((((.((.(((.((..((((..........)))).))))).)))))))...)))..)).))).)).)))))))))).))).)))&(((((((.((((.((((.((((((..((((((..........))))))..)))))).)))))))).)))))))'

pairing_conservation

UUUUUUUUUPPPUUPPPUPPPPPPPPPPPUUPPUUPPPPPUUUUPPPUUUPPPPPUPPUPPPUPPUUPPPPUUUUUUUUUPPPPUPPPPPUPPPPPPPPUUUPPPUUPPUPPPUPPUPPPPPPPPPPPUPPPUUPPP&PPPPPPPUPPPPUPPPPUPPPPPPPUUPPPPPPPUUUUUUUUUUPPPPPPPUUPPPPPPPUPPPPPPPPPPUPPPPPPP'

Legend: * = unknown, U = conserved unpaired, + = unpaired -> paired, x = unpaired -> pseudoknot P = conserved paired, p = paired -> pseudoknot,
- = paired -> unpaired S = conserved pseudoknot, s = pseudoknot -> paired, _ = pseudoknot -> unpaired

**C**

H19_Exon3='..............((((((....))(((((((((....)))))))).(((((......))))).((((.........)))).(((((....))))).)))))..........'
mir-675='(((((((.((((.((((.((((((..((((((..........))))))..)))))).)))))))).)))))))'
cofold='....((...((((((((((((((((((((((({....}))))))))..........(((.(((((....((((.(((..(((((((..((..((((,,(((............&....)))))))).)).)))))))))).))))))))).))).)))))).)))).)))),..,))))...)).'

pairing_conservation

'UUUU++UUU+++++PPPPPPP++++PPPPPPPPPpUUUUpPPPPPPPU-----UUU+++-PPPP+----++++U+++U-PPP+PPP--++UUPPPP**PPP--UUUUUUUUUU&----PPP+PPPPUPP--+PPPPPP++P-PPPP+++++U+++U-PPPPP+UPPPP-P+PP*--*PP+P---PP-'

Legend: * = unknown, U = conserved unpaired, + = unpaired -> paired, x = unpaired -> pseudoknot P = conserved paired, p = paired -> pseudoknot,
- = paired -> unpaired S = conserved pseudoknot, s = pseudoknot -> paired, _ = pseudoknot -> unpaired

**D**

H19_Exon4='.....((((((((((..((((......(((.((....((((((.((....(((...((.((....)).)).....(((((((..((.....((((....))))....))..))))).))..(((((.....)))))......(((((((...((((........))))))))))).................................)))....)).))))))....)).)))......)))))))))).......((((((((.((((((((((((((...((...((......(((.((((........)))))))......)).)).))))))))))))...))).)))))))))).(((..(((((...(((..(((((.......(((....))).((((((((((((((............))))))..((((((((..(((.(((.(((...)))..)))...)))..)))))))))((((((((((..........)))))).((......)).))))).............)))))))))..........((((((((((....)))))))))).)))))..)))....)))))..)))))))..'
miR-675='(((((((.((((.((((.((((((..((((((..........))))))..)))))).)))))))).)))))))'
cofold='.....((((((((((..((((......(((.((....((((((.((....(((...((.((....)).)).....(((((((..((.....((((....))))....))..))))).))..(((((.....)))))......(((((((...((((........))))))))))).................................)))....)).))))))....)).)))......)))))))))).......((((((((.((((((((((((((...((...((......(((.((((........)))))))......)).)).))))))))))))...))).)))))))))).(((..(((((...(((..(((((.......(((....))).((((((((((((((............))))))..((((((((..(((.(((.(((...)))..)))...)))..)))))))))((((((((((..........)))))).((......)).))))).............)))))))))..........((((((((((....)))))))))).)))))..)))....)))))..)))))))..&(((((((.((((.((((.((((((..((((((..........))))))..)))))).)))))))).)))))))'

pairing_conservation

'UUUUUPPPPPPPPPPUUPPPPUUUUUUPPPUPPUUUUPPPPPPUPPUUUUPPPUUUPPUPPUUUUPPUPPUUUUUPPPPPPPUUPPUUUUUPPPPUUUUPPPPUUUUPPUUPPPPPPUPPUUPPPPPUUUUPPPPPUUUUUUPPPPPPUUPPPPUUUUUUUUPPPPPPPPPPPPUUUUUUUUUUUUUUUUUUUUUUUUUUUUUUUUUUUPPUUUUPPUPPPPPPPUUUUPPUPPPPUUUUUPPPPPPPPPPPPUUUUUUPPPPPPPPPPUPPPPPPPPPPPPPPPPUUUPPUUUPPUUUUUUPPPUPPPPUUUUUUUUPPPPPPPPUUUUUPPUPPUPPPPPPPPPPPPPUUUPPPUPPPPPPPPPPUPPPUUPPPPPUUPPPUUPPPPPUUUUUUUPPUUUUPPPUPPPPPPPPPPPPPPPUUUUUUUUUUUPPPPPPUUPPPPPPPPPPUUPPPUPPPUPPPUUPPPUUPPPUUUPPPUUPPPPPPPPPPPPPPPPPPPPPUUUUUUUUUPPPPPPUPPUUUUUPPUPPPPPPUUUUUUUUUUUUUUPPPPPPPPPUUUUUUUUUPPPPPPPPPPUUUUPPPPPPPPPPPPUPPPPPPUUPPPUUUUPPPPPPUUPPPPPPPUU&PPPPPPPUPPPPUPPPPUPPPPPPPUUPPPPPPPUUUUUUUUUUPPPPPPPUUPPPPPPPUPPPPPPPPPPUPPPPPPP'

Legend: * = unknown, U = conserved unpaired, + = unpaired -> paired, x = unpaired -> pseudoknot P = conserved paired, p = paired -> pseudoknot,
- = paired -> unpaired S = conserved pseudoknot, s = pseudoknot -> paired, _ = pseudoknot -> unpaired
